# Identifying Genomic Alterations in Stage IV Breast Cancer Patients using MammaSeq™: An International Collaborative Study

**DOI:** 10.1101/2020.03.01.965632

**Authors:** Osama Shiraz Shah, Atilla Soran, Mustafa Sahin, Serdar Ugras, Esin Celik, Peter C. Lucas, Adrian V. Lee

## Abstract

**Background:** Identification of genomic alterations present in cancer patients may aid in cancer diagnosis and prognosis and may identify therapeutic targets. In this study, we aimed to identify clinically actionable variants present in stage IV breast cancer (BC) samples.

**Materials and Methods:** DNA was extracted from formalin fixed paraffin embedded (FFPE) samples of BC (n=41). DNA was sequenced using MammaSeq™, a BC specific next generation sequencing panel targeting 79 genes and 1369 mutations. Ion Torrent Suite 4.0 was used to make variant calls on the raw data and the resulting single nucleotide variants were annotated using CRAVAT toolkit. SNVs were filtered to remove common polymorphisms and somatic variants. CNVkit was employed to identify copy number variations. The Precision Medicine Knowledgebase (PMKB) and OncoKB Precision Oncology Database were used to associate clinical significance with the identified variants.

**Results:** A total of 41 Turkish BC patient samples were sequenced (read depth of 94 – 13340, median of 1529). These samples were from patients diagnosed with various BC subtypes including invasive ductal carcinoma (IDC), invasive lobular carcinoma (ILC), apocrine BC and micropapillary BC. In total, 59 different alterations (49 SNVs and 10 CNVs) were identified. From these, 8 alterations (3 CNVs – *ERBB2, FGFR1* and *AR* copy number gains and 5 SNVs – *IDH1*.R132H, *TP53*.E204*, *PI3KCA*.E545K, *PI3KCA*.H1047R and *PI3KCA*.R88Q) were identified to have some clinical significance by PMKB and OncoKB. Moreover, the top five genes with most SNVs included *PIK3CA, TP53, MAP3K1, ATM* and *NCOR1*. Additionally, copy number gains and losses were found in *ERBB2, GRB7, IGFR1, AR, FGFR1, MYC* and *IKBKB*, and *BRCA2, RUNX1* and *RB1* respectively.

**Conclusion:** We identified 59 unique alterations in 38 genes in 41 stage IV BC tissue samples using MammaSeq™. Ten of these alterations were found to have some clinical significance by OncoKB and PKMB. This study highlights the potential use of cancer specific NGS panels in clinic to get better insight into the patient-specific genomic alterations.

**Highlights:**

- 41 stage IV stage breast cancer patients of Turkish descent were sequenced using MammaSeq^™^
- 49 single nucleotide variations and 10 copy number variations identified
- *PIK3CA* and *TP53* mutations were present in 24% and 17% of the samples respectively
- 37% of the samples had *ERBB2*/*GRB7* gains and 7% had loss of *BRCA2*/*RB1* locus
- Eight clinically significant alterations were identified

**Micro Abstract:** We performed targeted sequencing using DNA from FFPE samples of 41 stage IV breast cancer patients using MammaSeq™, a breast cancer gene specific targeted sequencing panel. In total, 49 single nucleotide variations (SNVs) and 10 copy number variations (CNVs) were identified. Eight alterations (3 CNVs – *ERBB2, FGFR1* and *AR* copy number gains and 5 SNVs – *IDH1*.R132H, *TP53*.E204*, *PI3KCA*.E545K, *PI3KCA*.H1047R and *PI3KCA*.R88Q) were identified to have clinical significance by PMKB and OncoKB databases.

## INTRODUCTION

Next generation sequencing (NGS), with its improving speed, accuracy and cost, is becoming increasingly feasible in clinical care. However, current selection of therapies for breast cancer (BC) patients are primarily based on clinical and histological factors and genomic factors such as genomic heterogeneity are often overlooked^1^. Supplementing therapy selection with NGS diagnostics can help account for genomic heterogeneity and aid in designing effective treatment regimens for BC patients. NGS based diagnostics can be especially useful for management of stage IV BC disease which have a very heterogenous genetic landscape^2^. For such cases, NGS can identify which genomic features are/could be responsible for drug resistance and what alternate clinically actionable targets might be available.

In this study we investigated the genomic alteration landscape of stage IV BC patients of Turkish descent (Table 1) by sequencing formalin-fixed paraffin-embedded (FFPE) BC samples using MammaSeq™ ^3^. We identified 59 alterations of which 8 were clinically actionable. With this study we provide additional support for use of NGS diagnostics in management of stage IV BC patients.

**Table 1:**
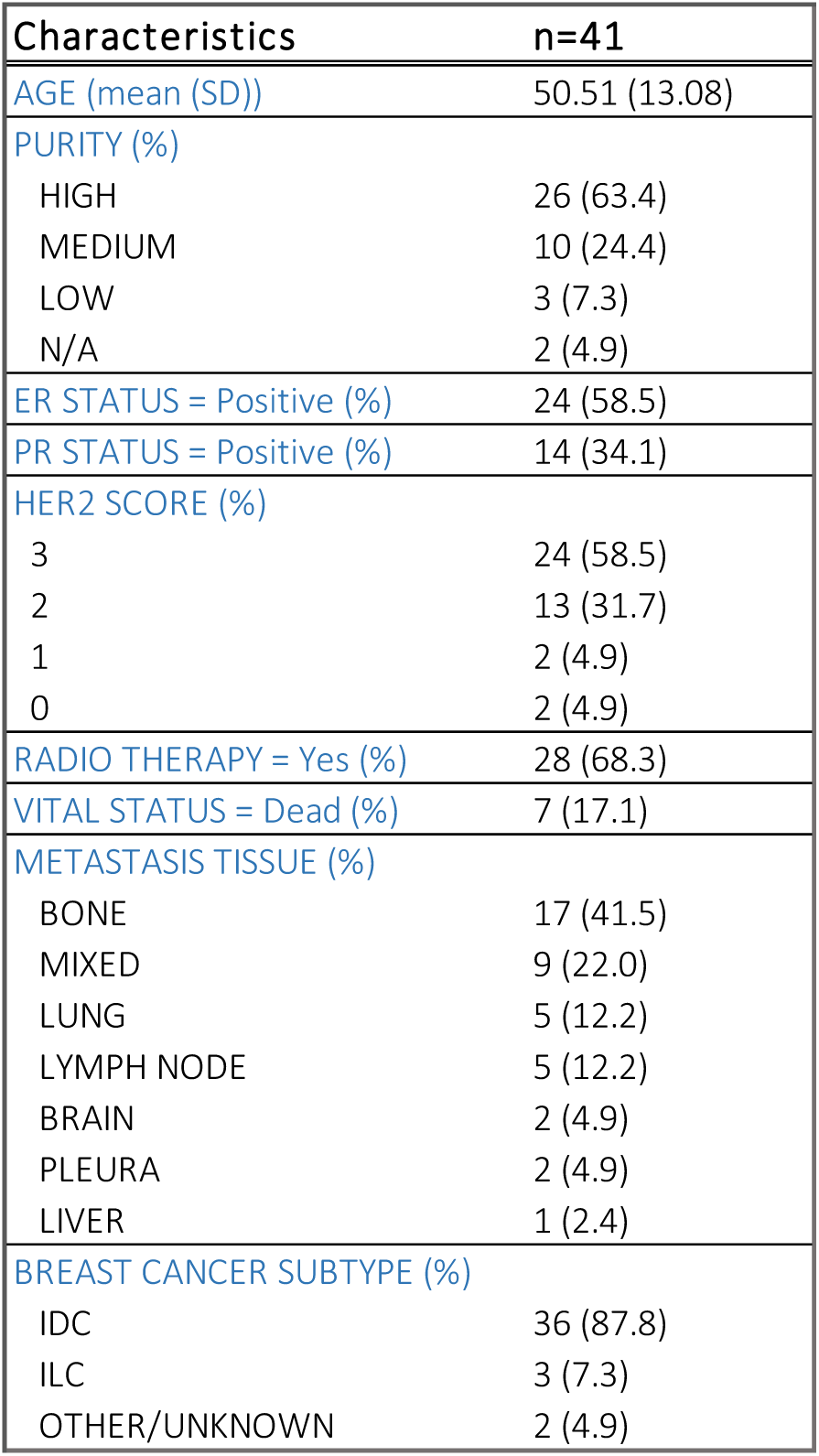
Patient and specimen characteristics.

| Characteristics | n=41 |
| --- | --- |
| AGE (mean (SD)) | 50.51 (13.08) |
| PURITY (%) |  |
| HIGH | 26 (63.4) |
| MEDIUM | 10 (24.4) |
| LOW | 3 (7.3) |
| N/A | 2 (4.9) |
| ER STATUS = Positive (%) | 24 (58.5) |
| PR STATUS = Positive (%) | 14 (34.1) |
| HER2 SCORE (%) |  |
| 3 | 24 (58.5) |
| 2 | 13 (31.7) |
| 1 | 2 (4.9) |
| 0 | 2 (4.9) |
| RADIO THERAPY = Yes (%) | 28 (68.3) |
| VITAL STATUS = Dead (%) | 7 (17.1) |
| METASTASIS TISSUE (%) |  |
| BONE | 17 (41.5) |
| MIXED | 9 (22.0) |
| LUNG | 5 (12.2) |
| LYMPH NODE | 5 (12.2) |
| BRAIN | 2 (4.9) |
| PLEURA | 2 (4.9) |
| LIVER | 1 (2.4) |
| BREAST CANCER SUBTYPE (%) |  |
| IDC | 36 (87.8) |
| ILC | 3 (7.3) |
| OTHER/UNKNOWN | 2 (4.9) |

## MATERIALS AND METHODS

### Patients and Samples Processing

Following review and approval by the Institutional Review Board at the University of Pittsburgh, we obtained FFPE tissues from BC cases diagnosed at Selcuk University, Konya, Turkey (Table 1, supplementary File 1). Samples were sectioned and reviewed by a breast pathologist at UPMC Magee Women’s Hospital (PCL). Sections were macro-dissected followed by deparaffinization (Qiagen cat#19093) and DNA isolation (QIAamp DNA FFPE kit, cat#56404), as per manufacturer’s instructions. Isolated DNA was quantified using the ThermoFisher Qubit dsDNA S/BR kit.

### Ion Torrent Sequencing

Samples with adequate DNA were sent for Ion Torrent Sequencing at the University of Pittsburgh Genomics Core following established methods^3^. In summary, we collected 20ng of DNA (divided equally between two pools of primers) for library preparation using Ion AmpliSeq™ library kit (ThermoFisher Scientific) and MammaSeq™ primer panel. Emulsion PCR was used for target enrichment using Ion OneTouch 2 system (ThermoFisher Scientific). Enriched library was loaded onto Ion chips and sequenced on Ion Torrent Personal Genome Machine (PGM™, ThermoFisher Scientific).

### Quality control analysis

TarSeqQC^4^, a R package for quality control analysis of targeted NGS experiments, was used for quality control analysis of the generated NGS dataset. Coverage of amplicons sequenced by primer mix 1 (pool 1) and primer mix 2 (pool2) were assessed.

### Variant calling

Variant calls were performed using Ion Torrent Suite 4.0^5^. Briefly, the raw fastq files were aligned to hg19 reference genome to generate VCF 4.0 files. The variants were prefiltered for allele frequency cutoff of 5%. For further variant filtering we followed established methods as described in Smith et al 2019^3^. Briefly, we annotated the variants using Cravat CHASM v4.3^6^ removed variations which were: 1) synonymous, 2) present in 30% or more of the samples, 3) having high strand bias, 4) found in gnomAD or 1000 genomes database with an allele frequency of more than 1%, 5) having allele frequency of more than 90% and 6) having a read depth of 50 or lower.

To compute CNVs, we used the Copy Number Ratio (CNR) outputs from CNVkit^7^. CNR files contain following: 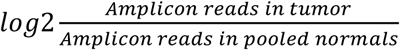 value for all gene amplicons. CNR value residing in each bin were aggregated and used to calculate a copy number value for each gene in that bin using the formula: 2 * 2^(*aggregated CNR*). CNR values with read depth lower than 5 and those that lied 2 standard deviations away from mean in each sample to discard low confidence amplicons and potential outliers. For detailed information on these calculations see CNVkit^7^ documentation (https://cnvkit.readthedocs.io/en/stable/pipeline.html).

### Clinical Actionability Analysis

We used PMKB^8^ and OncoKB to associate clinical significance to the shortlisted CNVs and SNVs. Briefly, we downloaded the databases and then cross checked the presence of each alteration in it.

### Data and code

R^9^ was used for quality control, variant annotation, filtering and clinical significance association as well as for making the barplots, venn diagram (VennDiagram^10^) and oncoprint (Complex Heatmaps^11^). The code is available openly at on github: https://github.com/osamashiraz/leeoesterreich_MammaSeq-Turkish.Cohort/.

## RESULTS AND DISCUSSION

### Cohort Overview and Sequencing

For this study, we obtained FFPE tissue blocks from BC cases diagnosed at Selcuk University, Konya, Turkey. The mean age of patients was 50.5 with standard deviation of 13.1 (Table 1). Tumor purity (“Purity”) values are based on the interpretations of the pathologist. ER, PR and HER2 status listed in Table 1 are based on Immuno-HistoChemistry (IHC) test. Most samples were ER+, PR- and HER2+. 41.5% of the samples were from bone metastasis tissue, 22% from mixed/complex tissue, 12.2% from lung and lymph node sites each, 5% from brain and pleura each and 2.4% from liver. Most of the samples (87.8%) were diagnosed as invasive ductal carcinoma (IDC), 7% were invasive lobular carcinoma (ILC) and few (5%) could not be diagnosed and were characterized as OTHER/UNKOWN (see Supplementary File 1 for details).

### Alteration Landscape

In total, all 41 samples were successfully sequenced using MammaSeq™ panel with mean coverage ranging from 94-133,340, average mean coverage of 2,308 and median of 1,529 (Supplementary Fig. 1). Variant calls were made using IonTorrent 4.0^5^ and copy number calls using CNVkit^7^. SNVs from variant calls were annotated using Cravat CHASM v4.3^6^ and were filtered to remove common polymorphisms and sequencing artifacts (see Supplementary Fig. 3) as described in Materials and Methods section. CNV computation was performed using CNR values from CNVkit^7^ as described in Materials and Methods. Gains were defined as copy number > 6 and losses as copy number < 1. During quality control, we noticed that amplicons sequenced by primer mix 2 (pool 2) had a lower overall coverage compared to primer mix 1 (pool 1) (Supplementary Fig. 2). To account for the influence of low coverage amplicons on variant identification and copy number computation, we removed low read depth variant calls from SNV analysis and low coverage amplicons from CNV analysis as mentioned in Materials and Methods section (also see Supplementary File 2). The final oncoprint showing the genomic alterations detected by MammaSeq™ is shown in Fig. 1.

**Figure 1:**
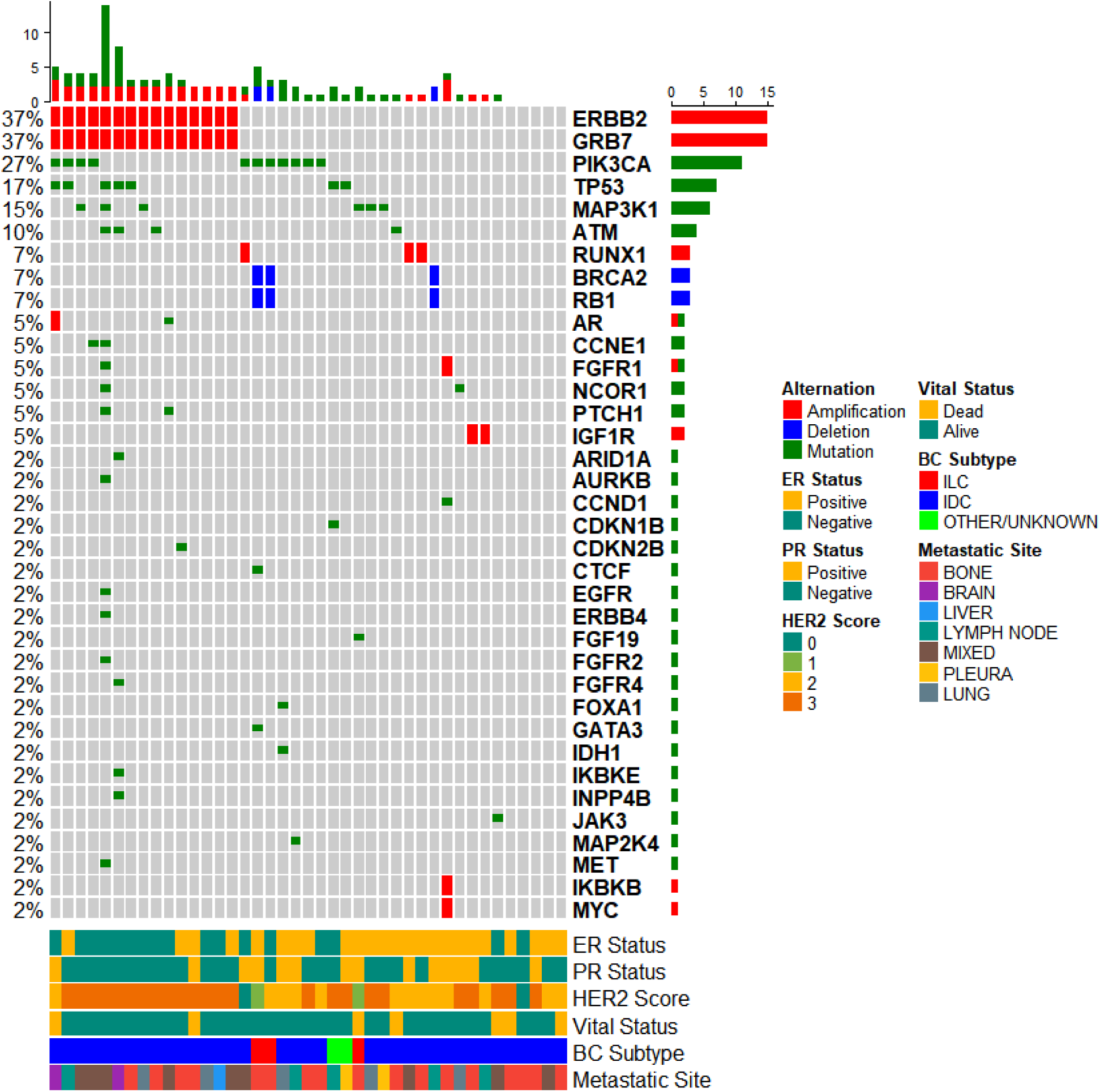
Oncoprint showing the genetic alterations identified by the MammaSeq™ gene panel in Turkish BC Cohort. On left the mutational frequency per gene for this cohort is reported. The corresponding genes are labelled on the right side. The barplots beside the gene labels report the mutational count. Bottom color bar represent the annotation based on clinical variables. The topmost barplots represent the mutation count per sample. IDC: Invasive ductal cancer and ILC: Invasive lobular cancer.

**Figure 2:**
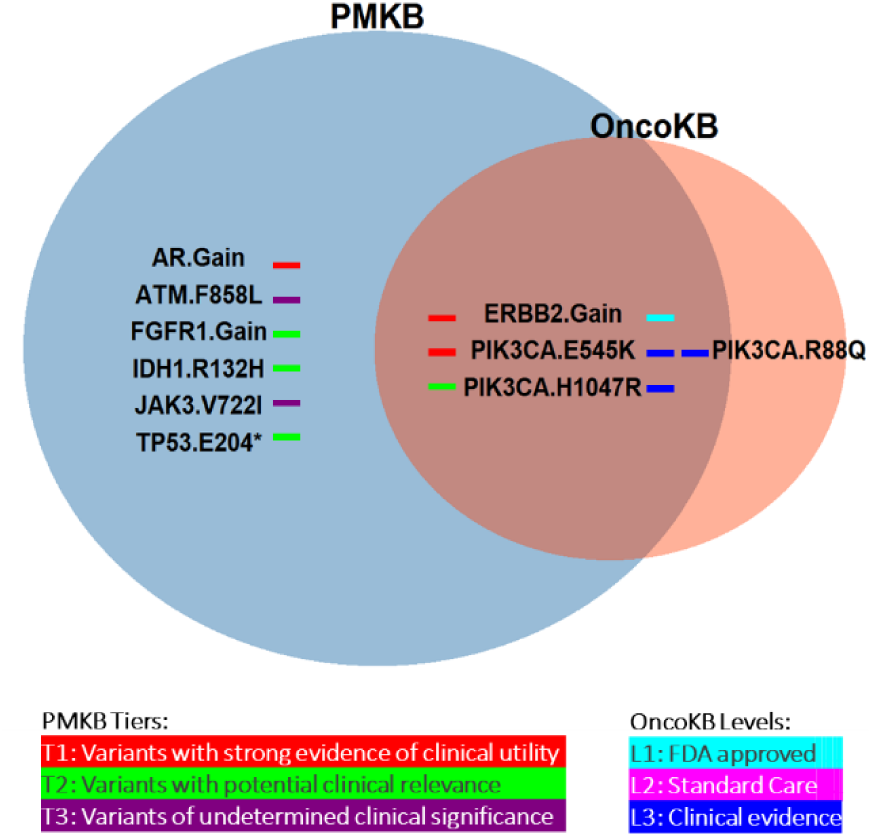
Overlap between interpretations from OncoKB and PMKB on top. Significance categories used by these databases on bottom.

We found a total of 59 unique alterations. Forty-nine of these were SNVs. *PIK3CA* gene is an important regulator of PI3K-AKT pathway which controls cell proliferation and survival^12^. In our analysis *PIK3CA* was the most frequently mutated gene. *PIK3CA* mutations were seen at rate of 27% in our cohort while other metastatic BC studies report 28%^13^, ∼29%^14^ and ∼30% (MBC project - The Metastatic Breast Cancer Project (https://www.mbcproject.org/), a project of Count Me In (https://joincountmein.org/). *PIK3CA* mutation rates seen in stage IV/metastatic BCs are lower than those reported by primary BC studies (∼35% in TCGA^15^ and 37% in METABRIC^16,17^). Regardless of these differences, *PIK3CA* mutations are widespread in primary and metastatic BCs. The FDA recently approved the first *PIK3CA* inhibitor, Alpelisib, for patients with *PIK3CA* mutation confirmed via a companion diagnostic^18^.

*TP53* is an important tumor suppressor which inhibits abnormal proliferation^19^. This was the second most frequently mutated gene in our cohort. The SNV rate in *TP53* was 17% while other metastatic BC studies have reported 13%^13^, ∼38%^14^ and 30% (MBC project). Both our and Pezo et al’s^13^ study use a targeted NGS approach; i.e. only hotspot regions of select genes were sequenced to identify SNVs. Targeted approaches are not comprehensive and could underrepresent the net rate of mutations, especially for tumor suppressor genes such as *TP53* where pathogenic mutations may be spread throughout the coding region. In contrast, the Lefebvre et al^14^ and MBC project use whole exome sequencing which offers more comprehensive SNV identification by covering all gene exons. This might explain why the latter two reports show higher numbers of *TP53* mutations. Furthermore, *TP53* gene mutation rate in primary BC studies was ∼34% ^15–17^ which overlaps with the rates seen in metastatic BCs.

*MAPK31* is MAPK kinase kinase protein that activates the MAPK/ERK pathway to control proliferation, migration and other cell fates^20^. We found *MAPK31* to be mutated at a rate of 15% compared to ∼8%^14^ and ∼4% (MBC project) in other metastatic BC studies. While in primary BC studies the mutation rate was ∼8%^15^ and ∼9%^14,16^.

*ATM* is a tumor suppressor which is important for DNA repair^21^. We observed the *ATM* gene to be mutated at a rate of 10%. Others have reported ∼4%^14^ and ∼2% (MBC project) for metastatic BCs. Apart from these frequently mutated genes we also found less frequent mutations in *AR, CCNE1, FGFR1, NCOR1, PTCH1, ARID1A, AURKB, CCND1, CDKN1B, CDKN2B, CTCF, EGFR, ERBB4, FGF19, FGFR2, FGFR4, FOXA1, GATA3, IDH1, IKBKE, INPP4B, JAK3, MAP2K4* and *MET* (see Fig. 1 and supplementary file 3). Oncoprint with frequency of each unique alteration is shown in Supplementary Fig. 4.

CNVs are an important class of genomic alterations. We identified 10 unique CNVs. Most common of these was *ERBB2*/*GRB7* locus gains. *ERBB2* is a receptor tyrosine kinase and belongs to the EGFR family. Aberrant activation of *ERBB2* is associated with more aggressive BCs and reduced patient survival^22^. We found *ERRB2*/*GRB7* gain in 37% of the cases compared to 22.36% in MBC project cohort. These values are higher than those reported for primary BCs^15–17^ (∼12-13%). Moreover, in our previous study we observed 2-fold higher expression of *ERBB2* gene in 35% of the brain metastases when compared to matched primary BC tissue^23^. These observations suggest that *ERBB2* gains may be frequently acquired as tumor progresses to metastatic disease. This makes it critical to examine the HER2 status of stage IV/metastatic patients to determine if HER2 therapies could be of therapeutic value.

We identified deletions of *BRCA2*/*RB1* locus at 7% rate (2/3 of these cases had ILC subtype). It is important to note that loss of *BRCA2* is associated with resistance to DNA damaging agents^24^. Similarly, *RB1* deficient tumors could show resistance to CDK4 inhibitors^14^. Hence, *RB1*/*BRCA2* locus genotyping in stage IV/metastatic BC patients can be of prognostic importance as it may help determine if cancer therapies such as CDK4/6 inhibitors would be effective or not.

*RUNX1* is an important cell fate regulator of ER+ mammary luminal cells and is often mutated in BCs^15,25^. *RUNX1* gains were seen in 7% of the samples. *RUNX1* amplifications are rare in both primary BCs^15–17^ (∼1-2%) and metastatic BCs^14^ (∼3%). Moreover, it’s role in BC tumorigenesis or metastasis has not been very well established. However, *RUNX1* upregulation in colorectal cancer has been reported to drive metastasis vis WNT pathway^26^. Further studies will be needed to validate our findings on *RUNX1* amplifications and their role in metastases.

*IGF1R* controls the insulin-like growth factor pathway, an important regulator of survival, motility and proliferation^27^. We also found *IGF1R* gains in 5% of the samples. This frequency was similar to that reported for primary BCs i.e. ∼3-5%^15–17^ and metastatic BCs ∼4%^14^. Based on these findings, *IGF1R* gains seem to have similar frequency in primary and metastatic BCs.

We also observed gains in *IKBKB, MYC, AR* and *FGFR1* at ∼2% frequency each. Except for *AR* gains, rest of the gene gains were reported at a much lower frequency by Lefebvre et al in metastatic BCs^14^ (*IKBKB*: ∼6%, *MYC*: ∼19%, and *FGFR1*: ∼13%). There could be few reasons for these differences. 1) As MammaSeq™ is a targeted panel and it does not cover the entire genes it targets which prevents comprehensive CNV discovery. 2) Also, the cohort is of small size and it does not capture the complete heterogeneity of stage IV/metastatic BC cases.

### Clinical Actionability of Identified Variant

Next, we sought to investigate the clinical actionability of alterations detected by MammaSeq™. For this purpose, we used two precision oncology databases; OncoKB^28^ and PMKB^8^. Eight alterations were highlighted by these databases (4 were reported only by PMKB, 1 only by OncoKB and 3 by both) (See Fig. 2). Variants of high clinical significance included *PIK3CA*-E545K mutation, *ERBB2* gain and *AR* gain. Among these, only *ERBB2* gain was commonly detected by both databases, while the other two were detected only by PMKB (Table 2). Other clinically actionable variants included: *FGFR1* gain (associated with endocrine resistace^29^), *IDH1*-R132H (associated with increased cell migration in gliomas^30^), *TP53*-E204*, *PIK3CA*-H1047R and *PIK3CA*-R88Q. All the identified *PI3K* mutations are activating in nature^31^. Among these, the most common alterations were *ERRB2* gain (37%), *PIK3CA*-H1047R (20%), and *PIK3CA*-E545K (7%) (see Supplementary Fig. 4). While remaining 5 alterations had a frequency of ∼2%.

**Table 2:** Clinically significant variants identified and annotated by OncoKB and PMKB along with information on allele frequencies and copy number changes.

| Sample ID | Gene | Alteration | Allele<br>Frequency | Copy<br>Number | Detected<br>by | PMKB<br>Tier | OncoKB<br>Level |
| --- | --- | --- | --- | --- | --- | --- | --- |
| lonXpress_001 | ERBB2 | Gain | - | 7 | Both | 1 | 1 |
| lonXpress_008 | ERBB2 | Gain | - | 41 | Both | 1 | 1 |
| lonXpress_010 | ERBB2 | Gain | - | 12 | Both | 1 | 1 |
| lonXpress_011 | ERBB2 | Gain | - | 13 | Both | 1 | 1 |
| lonXpress_012 | ERBB2 | Gain | - | 27 | Both | 1 | 1 |
| lonXpress_014 | ERBB2 | Gain | - | 20 | Both | 1 | 1 |
| lonXpress_017 | ERBB2 | Gain | - | 9 | Both | 1 | 1 |
| lonXpress_018 | ERBB2 | Gain | - | 11 | Both | 1 | 1 |
| lonXpress_031 | ERBB2 | Gain | - | 13 | Both | 1 | 1 |
| lonXpress_033 | ERBB2 | Gain | - | 29 | Both | 1 | 1 |
| lonXpress_036 | ERBB2 | Gain | - | 18 | Both | 1 | 1 |
| lonXpress_038 | ERBB2 | Gain | - | 8 | PMKB | 1 | 1 |
| lonXpress_039 | ERBB2 | Gain | - | 14 | PMKB | 1 | 1 |
| lonXpress_044 | ERBB2 | Gain | - | 7 | PMKB | 1 | 1 |
| lonXpress_045 | ERBB2 | Gain | - | 9 | Both | 1 | 1 |
| lonXpress_013 | PIK3CA | E545K | 0.79 | - | PMKB | 1 | 3 |
| lonXpress_015 | PIK3CA | E545K | 0.32 | - | Both | 1 | 3 |
| lonXpress_036 | PIK3CA | E545K | 0.41 | - | Both | 1 | 3 |
| lonXpress_001 | PIK3CA | H1047R | 0.41 | - | Both | 2 | 3 |
| lonXpress_010 | PIK3CA | H1047R | 0.31 | - | Both | 2 | 3 |
| lonXpress_011 | PIK3CA | H1047R | 0.50 | - | Both | 2 | 3 |
| lonXpress_020 | PIK3CA | H1047R | 0.31 | - | Both | 2 | 3 |
| lonXpress_021 | PIK3CA | H1047R | 0.30 | - | Both | 2 | 3 |
| lonXpress_029 | PIK3CA | H1047R | 0.21 | - | Both | 2 | 3 |
| lonXpress_037 | PIK3CA | H1047R | 0.36 | - | Both | 2 | 3 |
| lonXpress_043 | PIK3CA | H1047R | 0.49 | - | Both | 2 | 3 |
| lonXpress_037 | PIK3CA | R88Q | 0.35 | - | PMKB | - | 3 |
| lonXpress_001 | AR | Gain | - | 9 | Both | 1 | - |
| lonXpress_001 | TP53 | E204* | 0.54 | - | PMKB | 2 | - |
| lonXpress_046 | FGFR1 | Gain | - | 8 | Both | 2 | - |
| lonXpress_037 | IDH1 | R132H | 0.36 | - | Both | 2 | - |
| lonXpress_034 | ATM | F858L | 0.70 | - | Both | 3 | - |
| lonXpress_040 | JAK3 | V722I | 0.53 | - | OncoKB | 3 | - |

FDA recently approved the first *PIK3CA* inhibitor, Alpelisib, for treating *PI3KCA* mutant BCs^18^. *ERBB2* gains are seen in both primary BCs and metastatic BCs with a somewhat higher rate in the latter. Antibody-based, ERBB2-targeting therapeutics, including trastuzumab, have proven to be effective for these patients^32^. Thus, it is important to screen metastatic BC patients for *ERBB2* gains and provide treatment where applicable. Previously, it has been suggested that high *AR* expression has been associated with better response to endocrine-therapy in ER+ BCs^33^. However, recent data from BIG-1-98 did not report significant association of *AR* expression with prognosis or endocrine treatment differences^34^. Finally, FGFR family inhibitors for BCs are in testing and show promise^35^.

## CONCLUSION

With this study, we highlight the potential use of cancer specific NGS panels in diagnostic management of cancer patients in clinic. Such panels can be used in clinic to provide insight into the genomic alteration landscape of cancer patients and guide patient specific cancer therapy.^36^

## CLINICAL PRACTICE POINTS

Therapy selection for BC patients is primarily based on clinical and histological factors. However, genomic features such as genomic heterogeneity of BC play an in important role in predicting treatment response. NGS diagnostic assays are being widely developed and tested if they can improve conventional diagnostic and prognostic assays for different cancers. Such panels have been tested in BC. However, they have been limited to primary BC disease. We recently developed MammaSeq™ as an NGS based diagnostic/prognostic assay for both primary and metastatic BCs. In this study, we use MammaSeq™ to decode the underlying genomic heterogeneity of metastatic BC patients of Turkish descent and identify clinically actionable alterations.

We identified 59 unique alterations (49 SNVs and 10 CNVs) in 38 genes. *ERRB2* gains were the most frequent alteration and are targetable using anti-*HER2* drugs. Given the high frequency of *ERBB2* gains in metastatic BCs compared to primary BCs, it is critical to assess *HER2*/*ERBB2* status in metastatic BCs and treat accordingly. Moreover, various activating PI3K mutations were found. Patients harboring such mutations can benefit from Alpelisib (a recent FDA approved *PI3K* inhibitor). High rates of *MAP3K1* SNVs were identified, however, none of these have been reported as clinically actionable and further studies will be needed to evaluate their significance.

In summary, diagnostic panels such as MammaSeq™ can provide important genomic alteration information that can be used to evaluate BC patient disease heterogeneity and identify actionable drug targets.

## ACKNOWLEDGEMENTS

This project benefited from resources provided by the University of Pittsburgh HSCRF Genomics Research Core and the University of Pittsburgh Center for Research Computing. The authors would like to thank Fangping Mu for bioinformatics assistance. This work was supported by funding from Selçuk University.

