## Supplementary Data for "Identifying Genomic Alterations in Stage IV Breast Cancer Patients using MammaSeq™: An International Collaborative Study"

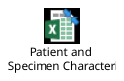

**Supplementary File 1**

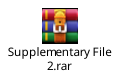

**Supplementary File 2**

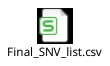

**Supplementary data 3**

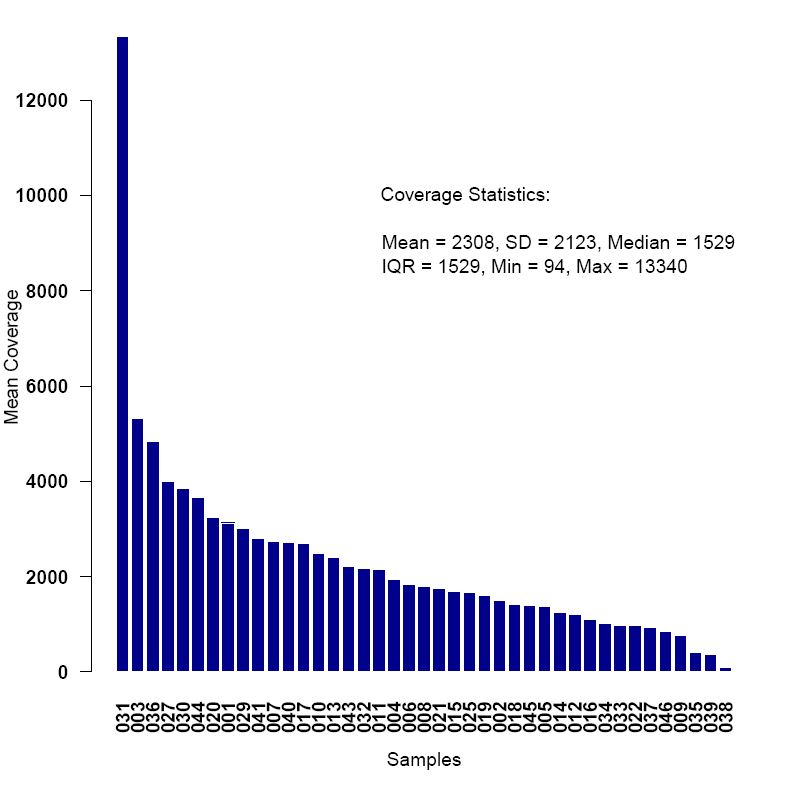

**Supplementary Figure 1: Mean Coverage across samples. The mean of mean coverages was 2308, standard deviation (SD) of 2123, median of 1529, Inter quartile range (IQR) of 1529, minimum coverage of 94 and max of 13340.**

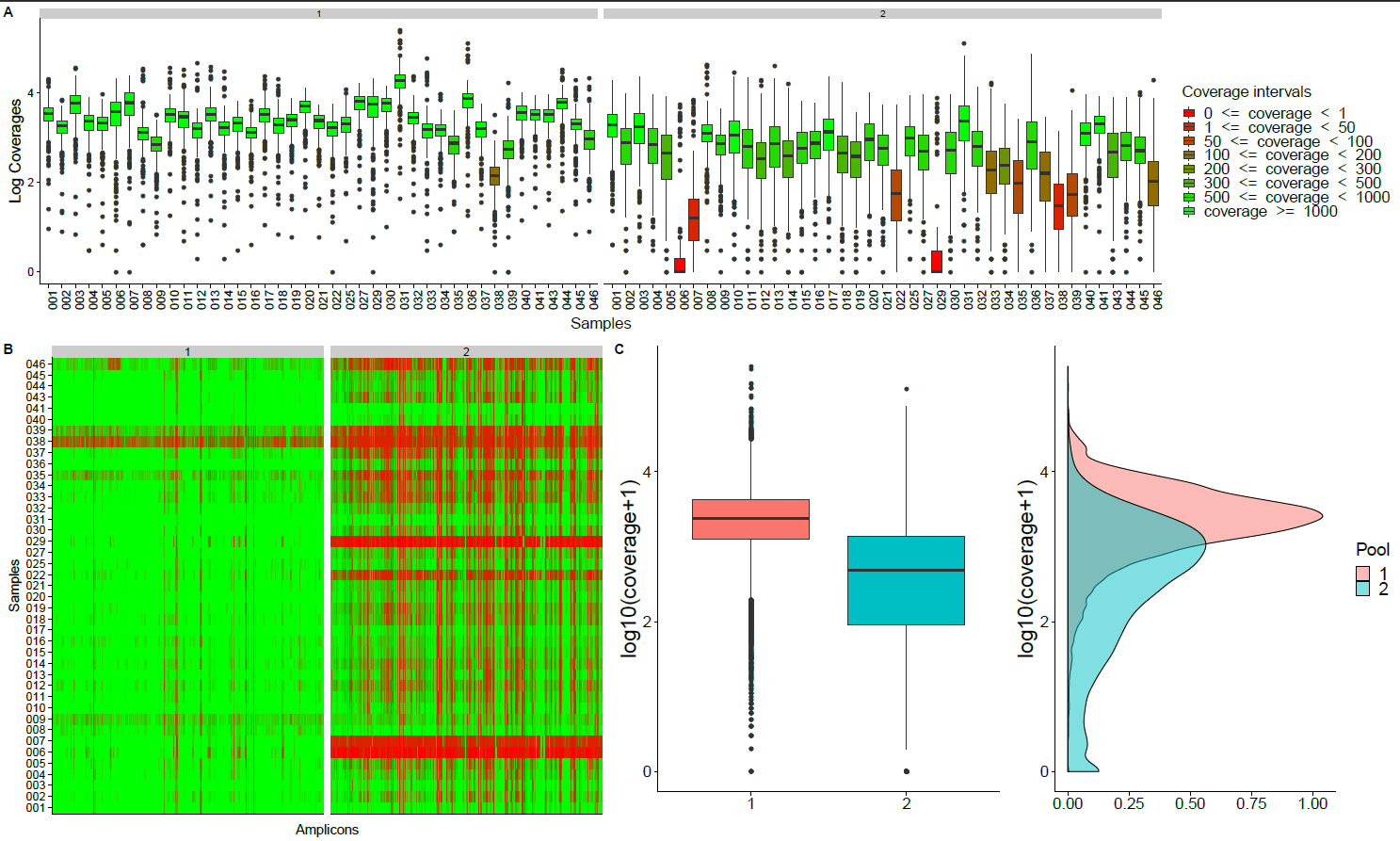

**Supplementary Figure 2: Quality control analysis. A) Log10 of overall coverage for pool 1 and pool 2 across samples. B) Amplicon coverages in each pool across sample. C) Comparison of distribution of log10 of coverages between pool 1 and 2.**

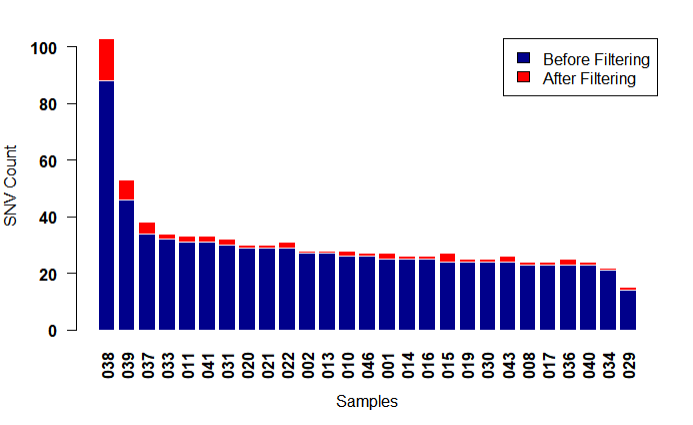

**Supplementary Figure 3: Single nucleotide variant (SNV) frequency before and after filtering. Filtering steps included: 1) removal of sequencing artifacts by removing variants found 30% or more of the samples, 2) selecting for high confidence variants by considering only the variants with strand bias within 2 standard deviations of the mean of the strand bias distribution i.e. between 0.45-0.58, 3) removing variants with reported allele frequency of more than 1% (common polymorphism) in 1000genome and gnomAD databases, 4) removing variants which were found with allele frequency of 90% or more in our sequenced samples, 5) removing synonymous variants and 6) removing variants with read depth lower than 50.**

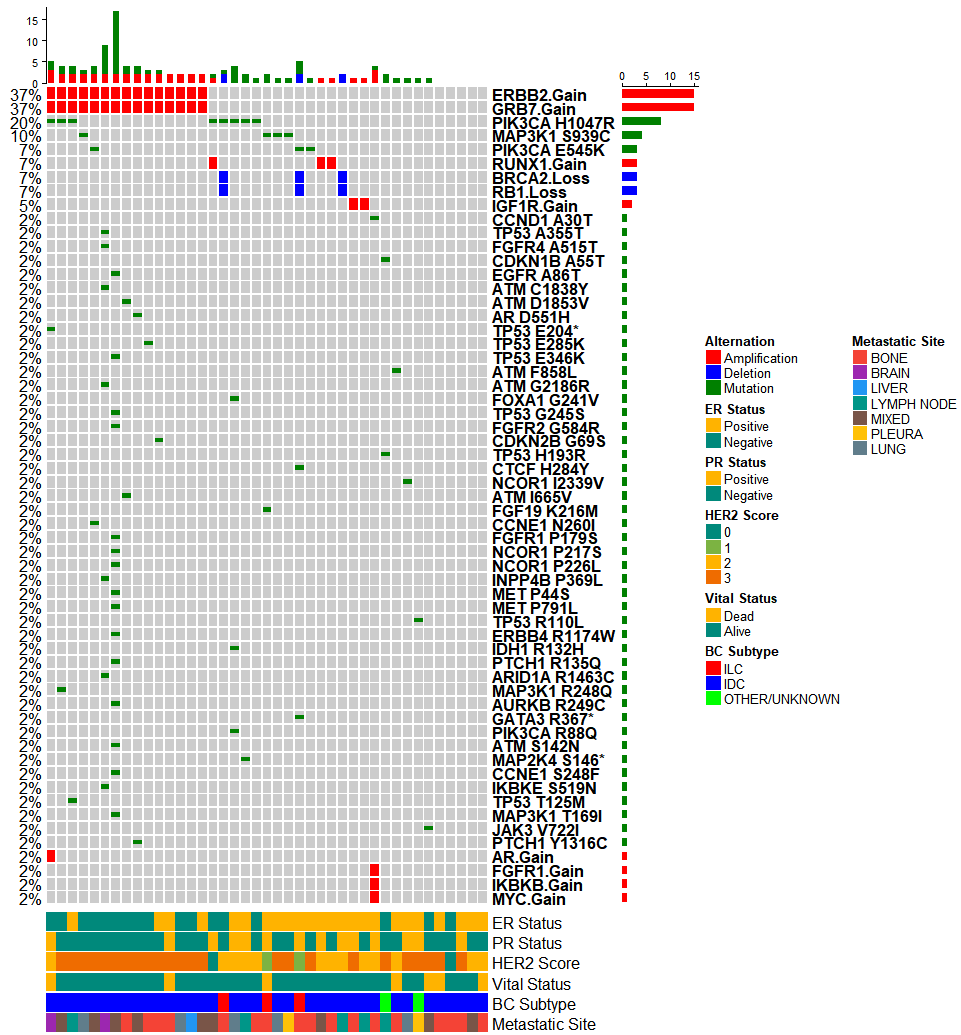

**Supplementary Figure 4: Oncoprint showing the genetic alterations identified by the MammaSeq^TM^ gene panel in Turkish BC Cohort.** **On left the mutational frequency per alteration for this cohort is reported. The corresponding alterations are labelled on the right side. The barplots beside the alteration labels report the mutational count. Bottom color bar represent the annotation based on clinical variables. The topmost barplots represent the mutation count per sample. IDC: Invasive ductal cancer and ILC: Invasive lobular cancer.**
